# Proteomic consequences of male mate preference learning and female phenotype in the butterfly *Bicyclus anynana*

**DOI:** 10.64898/2026.07.26.739888

**Authors:** Sushant Potdar, Dennis Province, Erica L. Westerman

## Abstract

Mate preference learning, where individuals use past social experience to choose mates, is prevalent in many species. Yet its consequences on reproductive investment, and whether individuals differentially invest in reproduction based on their learned perception of the mate’s phenotype, are unknown. We addressed these questions using the butterfly *Bicyclus anynana*, where males acquire individual preferences for artificially painted 0-UV dorsal hindwing spotted (DHSN) females, who then lay more eggs. We measured the spermatophore proteins transferred by naïve and experienced males to females manipulated to have preferred (0-DHSN) and unpreferred (2-DHSN) wing patterns. Using data independent acquisition (DIA), we identified 2144 proteins in the *B. anynana* spermatophore transferred to the female during mating. Experienced males that mated with preferred and unpreferred females had more differentially abundant (DA) proteins in their spermatophores that have functions in circadian rhythms, oogenesis, and neural signalling than naïve males. Proteins associated with oogenesis were DA in spermatophores transferred to preferred females, which may contribute to why preferred females lay more eggs. Overall, our study provides evidence for the role of experience-induced behavioural plasticity in tailoring male ejaculates in butterflies and identifies proteins that influence female physiology and behaviours which directly affect the pairs’ overall reproductive fitness.

## Introduction

Mate choice is a process where individuals choose their mates based on their preferences. However, these preferences can be innate or plastic [1]. One way preferences can be acquired or changed (plastic) is through mate-preference learning [2]. Mate-preference learning, where individuals acquire or change their preferences based on past social experiences, is prevalent in animals [2,3] and can lead to macroevolutionary changes, such as the evolution of novel behavioural traits and reproductive isolation [2–6]. While many studies focus on how mate-preference learning influences mate selection (pre-mating choice), relatively little is known about how mate preference learning influences behaviours exhibited during or after mating (peri- and post-mating choice).

Peri- and post-mating choice can be influenced by individual social environments through a myriad of morphological, physiological, and behavioural tactics employed by both males and females [1,7,8]. For instance, female soldier flies (*Merosargus cingulatus*) lay eggs immediately after copulating with males that perform female preferred copulatory courtship [9], whereas males copulate longer in the presence of male competition in the environment [10]. In *Drosophila melanogaster*, males copulate longer with larger females [11]. In cockroaches (*Nauphoeta cinerea*), males developed in social isolation transfer smaller spermatophores compared to group-reared individuals [12], whereas male mice (*Mus musculus*) secrete, and invest varied concentrations of reproductive proteins to females, based on their social status [13]. Thus, both males and females are able to alter their behaviour and reproductive investment based on their pre-mating preference and the social environment.

Males of many species transfer costly ejaculates to their partners during mating [14]. Since these ejaculates are often costly to produce, males may alter the size and content of their ejaculates based on their social environment and the perceived attractiveness of their mates [12,15]. In the context of sperm competition, males tend to give larger spermatophores to females [12,15–19], and change their ejaculate composition [20,21], as number of male competitors increase. While in the context of choice, males can change their ejaculate size and composition based on their perception of the female phenotype [15,22,23].

In insects, males, through their ejaculates, can influence many female post-mating behavioural and physiological changes [24,25] such as female aggression [26], feeding and dietary preferences [26–28], egg laying [29], female mate searching behaviour [30], female receptivity [14,18,31], sleep and circadian rhythm [32,33], and long-term memory formation [34]. These changes in female post-mating physiology and behaviour can directly influence male’s reproductive fitness [25]. To that end, many studies have characterized insect sperm and seminal fluid proteomes [35–48], but how mate preference learning —which can be a force in macro-evolutionary processes like assortative mating, reproductive isolation, and speciation [2–6]—affects the sperm and seminal fluid proteome is currently unknown. The proteome can also inform possible functions of these reproductive proteins, which may be shaped by the unique ecology of the said species. Taken together, it is imperative to understand and characterize spermatophore proteins, their influence on female physiology and behaviour, and the similarities and differences of the ejaculate proteomes between species.

Here, we use a butterfly system to assess whether males’ past social experience (mate preference learning), and males’ perception of the variation in female phenotype (based on his preference) affects his spermatophore investment to females during mating. In a number of species of butterflies, both males and females exert choice based on their preferences (mutual mate choice) for visual (wing colours and patterns) and odour cues (pheromones and anti-aphrodisiacs) [49–52]. During mating, male butterflies transfer a spermatophore to the female, which can be costly as it accounts for ∼3-23% of male body mass in some species [53,54]. Spermatophores can act as nuptial gifts by providing nutrients for female somatic maintenance and egg production [14,55,56], but can also act as a mating plug that reduces female receptivity [14,31,57]. However, we do not yet know if and how butterflies adjust their spermatophore composition based on their pre-mating preference and variation in female phenotype. In this study, we take advantage of a butterfly system, the species *Bicyclus anynana,* in which males acquire a preference for a female visual cue (number of UV-spots) through mate preference learning [58]. We use a spermatophore proteomics approach to qualify and quantify the spermatophore proteins transferred by naïve and experienced (trained) males to females with their preferred and unpreferred female wing pattern.

In *B. anynana,* individuals choose mates based on visual cues (UV-reflected spots on wings), and odour (pheromones) [51,52]. The wet season *B. anynana* males do not have an innate mating preference for the number of UV-reflected spots on the dorsal hindwing surface of females (0 DHSN vs 2 DHSN), but when sexually immature males just after emergence from their chrysalis are exposed to a 0-UV dorsal hindwing spotted (0 DHSN) female, they later prefer to mate with 0 DHSN females (mate preference learning) [58]. However, when newly emerged males are exposed to 2 DHSN females, they do not acquire a preference for 2 DHSN over 0 DHSN females, suggesting that *B. anynana* males are biased to learn preferences for reduced sexual ornaments [58]. Males’ perception of female phenotype (0 vs 2 DHSN) also appears to influence the spermatophore the male transfers to the female during copulation, as 2 DHSN females artificially modified to appear to have 0 DHSN lay more eggs after mating with a virgin male than females with their 2 DHSN visible to their male partner [59]. However, this difference in egg number is not associated with a significant difference in weight of spermatophore transferred to said female [59].

Since 0 DHSN females laid more eggs compared to 2 DHSN females without any significant changes in the male spermatophore weight, we hypothesized that males may be manipulating the protein contents of their spermatophore based on their perception of the female phenotype. We also hypothesized that males would change their spermatophore protein contents based on their previous social experience (naïve vs experienced with 0 DHSN female). To test these hypotheses, we conducted mating experiments in *B. anynana* to assess the effects of male social experience and males’ perception of female phenotype on various mating related behaviours (mating frequencies, the males’ latency to court, the pairs’ latency to mate, mating durations, and the spermatophore size), and the spermatophore protein content and quantity. We also describe the male transferred spermatophore proteome of the wet season *B. anynana* reared in greenhouse conditions, and compare the *B. anynana* spermatophore proteome to the sperm and seminal fluid proteins identified in other species. Our findings are important in understanding the realized reproductive potential of males based on their past social experiences, as well as their cryptic preferences.

## Materials and Methods

### 1. Study species and butterfly husbandry

The squinting bush brown butterfly *Bicyclus anynana* is found in the subtropical regions of East Africa, ranging from the equator to South Africa [60]. The Fayetteville AR greenhouse population is derived from a greenhouse bred population initiated from 80 gravid females collected from Malawi in 1988, and was started in 2017 from ∼1,000 eggs from a population in Singapore [61,62]. The population in Fayetteville AR is maintained in large colony cages (100 cm × 160 cm or 25.4 cm × 50.8 cm) where butterflies can mate freely and eggs are collected every week on corn plants. The caterpillars are reared to adulthood in subtropical wet season conditions by maintaining our greenhouse at ∼27°C and 70-80% RH with a 13:11 light:dark cycle. In these environmental conditions, the developing caterpillars will have wet season phenotype [63]. The caterpillars were fed with fresh corn (*Zea mays*) leaves *ad libitum* until pupation and were placed in cube cages (31.8 X 31.8 X 31.8 cm; Bioquip) until eclosion. Upon eclosion, the adult butterflies were transferred to sex- and age-specific cube cages and fed with banana, which was replaced every other day.

### 2. Mating assays

Mating assays took place from August 2022 till April 2023. All mating assays started 2 hours after dawn on the mating assay day, and ended when the pair mated or if there was no mating for 30 hours, and all assays used 2-day old males and females. On the day of emergence (day 0), males used in the mating assays were visually isolated in their respective rectangular mating assay cages (39.9 cm × 39.9 cm × 59.9 cm; Bioquip) with a wet banana as food for two days. To generate female 0 and 2 DHSN phenotypes, we used 1-day-old females that naturally possessed 2 DHSN on their wings and painted black paint (Testors enamel gloss black 1147) on top of the two UV reflective spots to make them 0 DHSN females. For 2 DHSN females, we painted the black paint just below the two naturally occurring UV reflective spots to remove the confounding effects of the presence of the paint (Figure S1) [58,59]. These females were then visually isolated from all males and females, in a separate cube cage (31.8 X 31.8 X 31.8 cm; Bioquip) for ∼24 hours with a wet banana for food and used in mating assays the next day.

For the mating assays, we used a full factorial treatment design using male experience (naïve vs experienced with 0 DHSN female) and female phenotype (0 vs 2 DHSN females) as factors, which created four treatments: 1. naïve male mated to 0 DHSN female (N0sp); 2. naïve male mated to 2 DHSN female (N2sp); 3. experienced male mated to 0 DHSN female (E0sp); 4. experienced male mated to 2 DHSN female (E2sp), which were the same treatments used in the previous *B. anynana* male mate-preference learning studies (Figure S2) [58,59]. For naïve treatments, on day 2, the isolated males were given either a 0 or 2 DHSN females to mate. For experienced treatments, on day 0, the newly emerged and visually isolated males were given a 3-hour exposure with a 2-day-old 0 DHSN female. Newly emerged males are sexually immature and will not mate during the 3-hour exposure period. After 3 hours, the 0 DHSN female is removed from the cage and the male is left alone for two days in his experimental rectangular cage. On day 3 (when the male is 2 days-old), the now experienced male is either given a 0 DHSN or a 2 DHSN female to mate. In each mating assay, we provided the pair with a fresh wet banana as a food and water source. We performed a total of 103 mating assays (Naïve paired with 0 DHSN female (N0sp) = 28; Naïve paired with 2 DHSN female (N2sp) = 28; Experienced paired with 0 DHSN female (E0sp) = 22; Experienced paired with 2 DHSN female (E2sp) = 25) to acquire the required number of spermatophores for downstream proteomic analyses (see below). During all the mating assays, we observed and recorded the time males took to court (latency to court), the time the pair took to mate (latency to mate), and the mating duration. Immediately after the copulation terminated, both the male and the female were sacrificed by freezing them in -30°C freezer, so that the females could not process and manipulate the spermatophore transferred by the male during mating. The mated butterflies were then transported to the lab for female abdomen dissection to extract the male transferred spermatophore. After the spermatophores were extracted from the female abdomen, we measured the butterfly’s forewing length from the base of the forewing to the apex using digital callipers as a proxy for male and female size. For this, we used all the mated pairs, as well as some unmated pairs (total N=76 pairs).

### 3. Spermatophore extraction

After every successful mating, the butterflies were transported to the lab after sacrificing them by freezing in -30°C. The female abdomen was dissected in 1X Phosphate-Buffered Saline (PBS) solution and the spermatophore with the female bursa copulatrix was extracted into the PBS solution using Zeiss stemi 508 stereo microscope. The spermatophore was then washed in fresh 1X PBS solution to remove female reproductive tract materials (like fat bodies and eggs) and photographed using Zeiss axiocam 305 colour camera. We stored the spermatophores within the female bursa copulatrix as we noticed that there were probable male-transferred proteins outside the spermatophore, but within the bursae, which we did not want to discard (Figure 1D). Hence, the proteomics were performed on the spermatophores and the accessory substances including the female bursa copulatrix, and the results should be interpreted cautiously as probable female derived proteins might be present in the spermatophore proteomic descriptions. The spermatophores were stored in 1X PBS solution at -20°C before shipping them to the IDeA National Resource for Quantitative Proteomics facility, Little Rock, AR for proteomic analyses. The photographed spermatophores were used to measure the area occupied by the spermatophore and used as a proxy for spermatophore size using *ImageJ* open access software as described in [64]. Briefly, we set the scale based on the scale of the original image and highlighted the spermatophores to separate it from the background (usually black). We then used the “wand” tracing tool to select the highlighted area and used ROI manager under the analyse and tools sections to add and measure the highlighted area which is the area occupied by the spermatophores. We visually assessed the end of the spermatophore tail inside the female bursa copulatrix and manually subtracted the area of the bursa not occupied by the spermatophore tail from the total spermatophore area.

**Figure 1:**
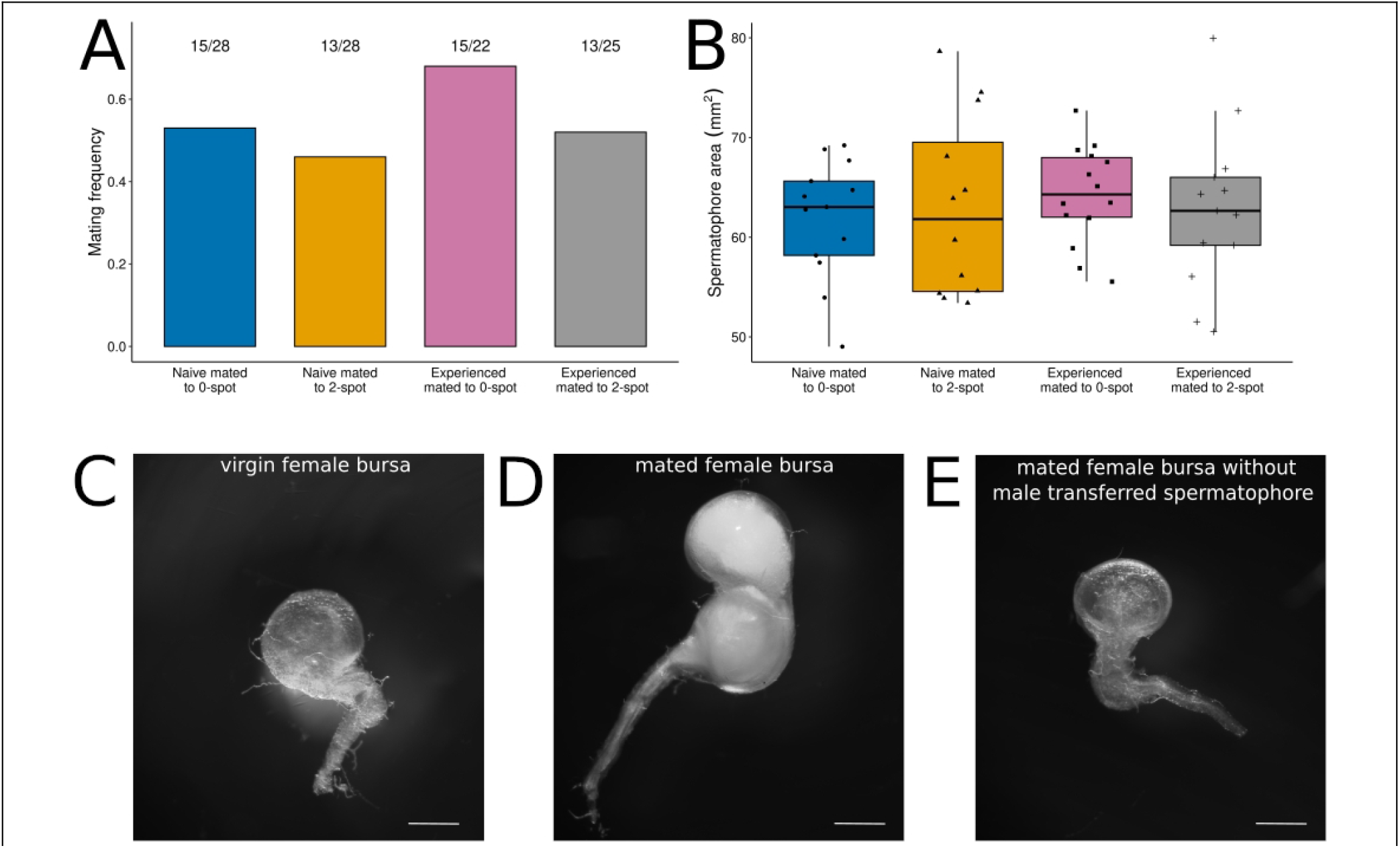
Mating frequency and spermatophore size does not change across treatments. A) The mating frequency of pairs do not change across treatments. The numbers at the top of each bar provides the number of matings over total number of trials in that treatment. B) The spermatophore area (measured as a proxy for size) does not change across treatments. C) A representation of a virgin female bursa copulatrix. D) A representation of a mated female bursa copulatrix with male transferred spermatophore and accessory substances. E) A representation of a mated female bursa copulatrix without a male transferred spermatophore and accessory substances. These bursae look similar to virgin female bursa copulatrix. The scale bar represents 100 µm

### 4. Spermatophore sample preparation

Based on a pilot run of *B. anynana* spermatophore proteomics at the IDeA National Resource for Quantitative Proteomics facility, we pooled two spermatophores from the same treatment to obtain the required quantity of proteins per sample. This provided us with N=6 samples per treatment (total of 24 samples with 48 spermatophores). The two spermatophores that were pooled were as similar as possible based on the pair’s latency to mate within each treatment. We used data independent acquisition (DIA) pipeline to qualify and quantify spermatophore proteins as it provides label-free and an unbiased analyses of peptides within all samples and identifies all the proteins present in the spermatophore above a certain mass to charge (m/z) ratio. After the samples were received at the proteomics facility, they were homogenized using a probe sonicator for 10 seconds at 10 Hz pulse rate. The samples were passed through a PBS washed Qia shredder filter by centrifuging for 5 minutes at 5000 x g. Proteins were quantified using Pierce BCA protein assay kit per manufacturer’s protocol. The BCA protein assay kit uses varied concentrations of bovine serum albumin (BSA) protein standards for colorimetric estimation of the protein samples. Later, we used 50µg of each sample and reduced them to a final concentration of 10mM using 5µL of 200mM TCEP and incubated it at 37°C for 30 minutes. We later alkylated the TCEP reduced samples to 30mM using 6.1µL of 500mM iodoacetamide solution for 60 minutes at room temperature in the dark. We then performed chloroform-methanol extractions to remove any contaminants like detergents that can block the LC-MS column [65]. The detergent free proteins, after air drying, were digested using trypsin (50:1 protein:trypsin ratio) in 100µL of 100mM TEAB [65]. After incubating this solution at 37°C for ∼20 hours, the proteins were desalted using a 96-well Sep-Pak (Waters Corporation) by repeated washing using a wash buffer (Buffer A = 0.1% formic acid + 0.5% acetonitrile) and finally eluting it with an elution buffer (Buffer B = 0.1% formic acid + 99.9% acetonitrile). The eluted samples were dried using a speed-vac and stored in -20°C until they were loaded onto the LC-MS. During the sample preparation process, we lost one sample from E0sp treatment, which left us with N=5 samples for that treatment only.

### 5. Liquid Chromatography tandem Mass Spectrometry (LC-MS)

The samples were serially loaded onto an Orbitrap Eclipse Mass Spectrometer (Thermo) by way of an UltiMate 3000 RSLCnano system (Thermo). Separation of the tryptic peptides were achieved by reverse phase XSelect CSH C18 2.5um resin (Waters) on an in-line 150 x 0.075 mm column. Peptides were eluted using a 60 min gradient from 98:2 to 65:35 buffer A:B ratio. Eluted peptides were ionized by electrospray (2.2 kV) followed by mass spectrometric analysis. To assemble a chromatogram library, six gas-phase fractions were acquired on the Orbitrap Eclipse with 4 m/z DIA spectra (4 m/z precursor isolation windows at 30,000 resolution, normalized AGC target 100%, maximum inject time 66 ms) using a staggered window pattern from narrow mass ranges using optimized window placements. Precursor spectra were acquired after each DIA duty cycle, spanning the m/z range of the gas-phase fraction (i.e. 496-602 m/z, 60,000 resolution, normalized AGC target 100%, maximum injection time 50 ms). For wide-window acquisitions, the Orbitrap Eclipse was configured to acquire a precursor scan (385-1015 m/z, 60,000 resolution, normalized AGC target 100%, maximum injection time 50 ms). This was followed by 50x 12 m/z DIA spectra (12 m/z precursor isolation windows at 15,000 resolution, normalized AGC target 100%, maximum injection time 33 ms) using a staggered window pattern with optimized window placements. Precursor spectra were acquired after each DIA duty cycle.

### 6. Statistical analyses of behaviour data

After extracting and measuring the behavioural and spermatophore data, we fit a generalized linear model (GLM) with mating (yes/no) as the response variable, and male experience (naïve, experienced), female phenotype (0 DHSN, 2 DHSN), and their interaction as the predictor variables using the binomial distribution. Later, we ran separate linear models with the behavioural data (latency to court, latency to mate, mating duration), and the area of the spermatophores as the response variables, and male experience (naïve, experienced), female phenotype (0 DHSN, 2 DHSN), and their interaction as the predictor variables. We also fit separate linear models with forewing length as the response variable and sex (male/female), followed by male experience, female phenotype, and their interactions as the predictor variables. All analyses and figures were generated using R software 4.4.1 [66].

### 7. Proteomics data analyses

Following data acquisition, we searched data using an empirically corrected library against the UniProt *Bicyclus_anynana* database (January 2023) and performed a quantitative analysis to obtain a comprehensive proteomic profile. Proteins were identified and quantified using EncyclopeDIA [67] and visualized with Scaffold DIA using 1% false discovery thresholds at both the protein and peptide level. Protein MS2 exclusive intensity values were assessed for quality using ProteiNorm [68]. The data were normalized using cyclic loess [69] and analysed using proteoDA to perform statistical analysis using Linear Models for Microarray Data (limma) with empirical Bayes (eBayes) smoothing to standard errors [69]. Proteins with an FDR adjusted p-value < 0.05 were considered significant. Proteins were identified and quantified using EncyclopeDIA, the proteome of *Bicyclus anynana* retrieved from UniProt (Taxon ID: 110368) and *Bicyclus anynana* genome retrieved from NCBIRefSeq (assembly: GCF_947172395.1) [70]. The MS2 spectral intensities of the identified proteins were normalized using cyclic loess and linear models for microarray (limma).

### 8. Functional annotation

The functional annotation for *B. anynana* was obtained from Ernst et al 2021 [62]. Briefly, the *B. anynana* genome was blasted against NCBI ‘nr’ database and the top 10 best blast hits at E-value <10^-3^ were obtained. The functional classifications were carried out using Blast2GO v5.2.5 [71] to identify the functional categories that each gene belongs to using InterProScan. The mapping and annotations were performed using default parameters and the final functional annotation table exported for downstream analyses.

### 9. GO enrichment analyses

We performed GO enrichment analyses using Fisher’s exact test, first for the spermatophore proteome, and then for the differentially abundant (DA) proteins from each treatment comparisons separately. For the GO enrichment analysis of the spermatophore proteome, we used the spermatophore proteins as the test set and the *B. anynana* genome as the reference set using FDR<0.01 for multiple testing in the “FilterMode” using Blast2GO 6.0.3 software [71]. We then reduced the obtained GO terms to most-specific terms using FDR<0.01. We compared the obtained *B. anynana* spermatophore GO terms with those from other lepidoptera spermatophores such as *Heliconius melpomene, Heliconius erato* [39,40], and the sperm proteomes of the monarch, *Danaus plexippus,* the cabbage white *Pieris rapae*, and tobacco hornworm, *Manduca sexta* [36,38,43], and with the sperm and seminal fluid proteome of the fruitfly *Drosophila melanogaster* [35,72]. Later, we separately ran Fisher’s exact test to identify GO terms with significantly DA proteins for each treatment comparison as the test set and the spermatophore proteome, and the *B. anynana* genome as the reference set using FDR<0.05 for multiple testing in the “FilterMode”. We performed GO enrichment analyses for the originally DA proteins as well as the permuted DA proteins (see below).

### 10. Sperm and seminal fluid proteins ortholog identification

We used OrthoFinder v 3.1.3 [73,74] to identify *Bicyclus anynana* spermatophore protein orthologs with other lepidopteran and non-lepidopteran species (*Aedes aegypti, Apis mellifera, Bombus terrestris, Drosophila melanogaster, Manduca sexta, Pieris rapae, Danaus plexippus, Heliconius erato, Heliconius melpomene, Mycalesis mineus,* and an outgroup bird species (*Taeniopygia guttata*)). We chose these representative species as their reproductive proteins (sperm, seminal fluid, or both in some cases) have been described (except *Mycalesis mineus*) [35–40,42–44,47,72,75]. We included *Mycalesis mineus* as it is the closest relative of *Bicyclus anynana* (diverged ∼25 mya) with an annotated genome [76]. We downloaded the latest reference proteomes of *Aedes aegypti, Apis mellifera, Bombus terrestris, Drosophila melanogaster,* and *Taeniopygia guttata* from NCBI, and the proteomes of *Heliconius erato, Heliconius melpomene, Pieris rapae* (HIRISE)*, Danaus plexippus,* and *Bicyclus anynana* from LepBase (https://lepbase.cog.sanger.ac.uk/#archive/v4/sequence/). *Manduca sexta* proteome (mansex_OGSv2.0) was downloaded from USDA (https://agdatacommons.nal.usda.gov/articles/dataset/Manduca_sexta_Official_Gene_Set_v2_0/24853296?file=44527493), and *Mycalesis mineus* proteome was downloaded from the Dryad repository from [76]. We extracted the longest transcript from these proteomes using scripts from OrthoFinder github, as well as custom scripts. 91.6% of genes were assigned to at least one orthogroups. We filtered and extracted 1:1 orthologs between *B. anynana* spermatophore proteins and compared it to the sperm and seminal fluid proteins described from other species.

### 11. Signal Peptide identification

We used SignalP 6.0 [77] to identify potentially secreted proteins in the *B. anynana* spermatophore. It identifies a predicted signal peptide sequence in the proteins present in the spermatophore. However, this is a conservative approach as other possible pathways can produce secretory proteins that may not be identified using SignalP 6.0 [78].

### 12. Permutation test

After normalization of the MS2 spectral intensities, we ran permutation tests to identify whether the observed differences in normalized mean protein intensities between treatments were different from the differences in simulated mean protein intensities between those same treatments. The null hypothesis of the permutation test is that the difference in observed and simulated means of protein intensities between two treatments are not different. We ran a 1000 permutation simulation for each protein within a specific two-treatment comparison and generated permuted p-values which were used to identify whether the specific protein was differentially abundant (DA) between the treatments. We performed these permutation tests for six treatment comparisons: 1. E0sp vs E2sp; 2. E0sp vs N0sp; 3. E2sp vs N2sp; 4. N0sp vs N2sp; 5. Experienced vs naïve; 6. 0 DHSN vs 2 DHSN.

## Results

### 1. Mating frequency, latency to court, latency to mate, mating duration, and spermatophore area does not change across treatments

The mating frequency in our one-choice assays did not change with male experience (naïve/experienced with 0 DHSN female; OR=0.53, CI=0.16-1.7, p=0.3), female phenotype (OR=0.51, CI=0.14-1.64, p=0.26), and their interaction (OR=1.5, CI=0.3-7.41, p=0.62; Figure 1A, Table S1), however, there was a trend where males paired with 0 DHSN females did mate a greater number of times (60%) in fewer mating trials compared to males paired with 2 DHSN females (49%). Male experience, female phenotype, and their interaction did not change the males’ latency to court the female (Male experience: t=0.66, p=0.51; Female phenotype: t=-0.65, p=0.51, Male experience*Female phenotype: t=0.54, p=0.59; Figure S3A, Table S1), the pairs’ latency to mate (Male experience: t=0.35, p=0.72; Female phenotype: t=0.88, p=0.37; Male experience*Female phenotype: t=-0.08, p=0.93; Figure S3B, C, Table S1), the pairs’ mating duration (Male experience: t=0.46, p=0.64; Female phenotype: t=-0.33, p=0.73; Male experience*Female phenotype: t=0.67, p=0.5;Figure S3D, Table S1), or the area of the spermatophores (proxy for size) transferred by males (Male experience: t=-0.83, p=0.4; Female phenotype: t=-1.07, p=0.28; Male experience*Female phenotype: t=0.17, t=0.86;, Figure 1B, Table S1). Based on the males’ and females’ forewing measurements, while males are smaller than females (t=-16.5, p=<2e-16), their size did not differ with respect to the male experience, female phenotype, and their interaction (Male experience: t=-0.59, p=0.55; Female phenotype: t=-0.39, p=0.69; Male experience*Female phenotype: t=0.4, p=0.68; Figure S3E, F, Table S1), Thus, the downstream spermatophore proteomic differences are unlikely to be due to simple differences in male or female size between treatments.

While it is generally assumed in butterflies that males always transfer spermatophores to females when they copulate (at least when those copulations are not interrupted) (Figure 1D), we found that for three pairs that mated and were not interrupted, the female bursa copulatrix did not contain any spermatophores or male secreted accessory substances (Figure 1E), and looked similar to the bursa of the virgin female (Figure 1C). These three pairs belonged to naïve male treatments and two of these had the highest mating durations observed in the experiment (47 and 100 minutes).

### 2. Spermatophore proteome

We recovered 2144 spermatophore proteins (Table S2) which is comparable to the sperm and seminal fluid proteomes of other species [35–40,42–44,47,72,75]. The GO enrichment analyses of the spermatophore proteome recovered 439 significantly enriched GO terms (FDR<0.01; Table S3) with 92 belonging to cellular components (CC, Figure S4A), 122 belonging to molecular functions (MF, Figure S4B), and 225 belonging to biological processes (BP, Figure S4C). When reduced to the most specific GO terms (FDR<0.01), we recovered 102 significantly enriched GO terms with 18 belonging to CC, 48 belonging to MF, and 36 belonging to BP (Table S4). The top most-specific enriched GO terms in the spermatophore proteome include “extracellular space”, “actin cytoskeleton”, and “microtubule” in the CC category; “ATP binding”, “GTP binding”, and “structural constituent of ribosome” in the MF category; and “protein folding”, “intracellular protein transport”, and “microtubule-based processes” in the BP category (Figure 2A).

**Figure 2:**
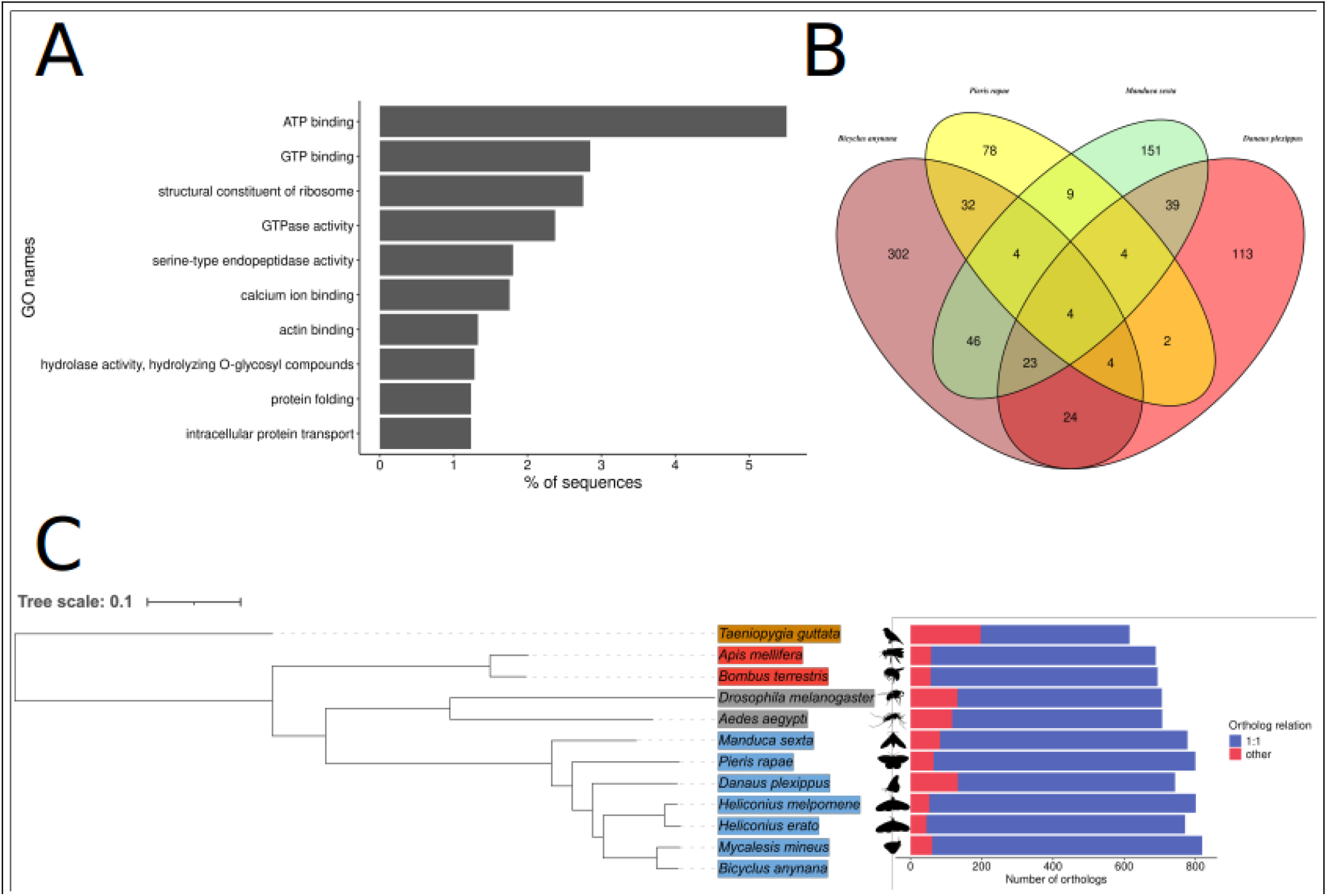
*Bicyclus anynana* spermatophore proteome. A) The top most-specific enriched GO terms of the proteins present in the male transferred spermatophore, with higher percentage of binding proteins. B) 44 GO terms were common between the spermatophore proteome of *B. anynana* and the sperm proteome of *Pieris rapae,* 55 GO terms were common between the spermatophore proteome of *B. anynana* and the sperm proteome of *Danaus plexippus*, and 77 GO terms were common between the spermatophore proteome of *Bicyclus anynana* and the sperm proteome of *Manduca sexta*. C) Greater number of *B. anynana* spermatophore orthologs were identified in closely related species. Species highlighted in blue are Lepidoptera (butterflies and moths), those in grey are Diptera (true flies), those in red belong to Hymenoptera (bees), with an outgroup bird species highlighted in yellow. In the bar plot, blue bars represent 1:1 orthology with *B. anynana* spermatophore proteins, where as red represents other relationship (1:many; many:1; many:many). Silhouettes were obtained from phylopic (phylopic.org)

We compared the *B. anynana* spermatophore enriched GO terms with the GO terms from the spermatophore proteome of *Heliconius melpomene*, and *H. erato*; and with the sperm proteome of *Danaus plexippus, Manduca sexta,* and *Pieris rapae,* and the sperm and seminal fluid proteome of *Drosophila melanogaster.* We identified 70 GO terms common with *H. melpomene* (Table S5), 74 GO terms common with *H. erato* (Table S5), 55 GO terms common with *D. plexippus* sperm proteome (Figure 2B, Table S5), 44 GO terms common with *Pieris rapae* sperm proteome (Figure 2B, Table S5), 77 GO terms common with *M. sexta* sperm proteome (Figure 2B, Table S5), and 32 GO BP terms common with the sperm and seminal fluid proteome of *Drosophila melanogaster* (Table S6). These terms were different based on which proteome we were comparing our *B. anynana* spermatophore proteome. Common GO terms between *B. anynana* spermatophore proteome and the *Heliconius* spermatophore proteomes included many “binding” terms (RNA binding, protein binding, calcium ion binding, metal ion binding) whereas when compared to the *Danaus plexippus* and *Manduca sexta* sperm proteome, we identified only two “binding” terms (ion binding, and purine ribonucleoside triphosphate binding), and “RNA binding” and “protein binding” with *Pieris rapae* sperm proteome. We retrieved many purine associated GO terms common with the *D. plexippus* and *M. sexta* sperm proteome. Looking at the common biological processes (BP) GO terms between *B. anynana* spermatophore proteome, and *D. melanogaster* sperm and seminal fluid proteome revealed many ATP associated processes such as “ATP metabolic processes”, and “ATP synthesis coupled electron transport”.

### 3. B. anynana spermatophore orthologs decrease with phylogenetic distance

Using *OrthoFinder 3.1.3*, we assigned 91.6% of genes to one or more orthogroups across the 12 species (see methods). After filtering the orthogroups for the spermatophore proteins of *B. anynana*, we found between 615 *B. anynana* spermatophore protein orthologs in *Taeniopygia guttata* (distantly related to *B. anynana*, Figure 2C) and 819 *B. anynana* spermatophore protein orthologs in *Mycalensis mineus* (most closely related to *B. anynana*, Figure 2C). Of these orthologs, 68%-94% had a 1:1 relationship with that of the *B. anynana* spermatophore proteins, with the highest percentage shared with closely related butterflies (*H. erato, H. melpomene,* and *M. minues*, Figure 2C), while the lowest percentage shared with distantly related species (*Taeniopygia guttata*, Figure 2C).

When we extracted the orthologs from the sperm proteomes of *Danaus plexippus*, *Pieris rapae,* and *Manduca sexta,* we identified the least number of *B. anynana* spermatophore orthologs with the sperm proteome of *D. plexippus* (73; Table S7), followed by *M. sexta* (147; Table S7), and *P. rapae* (263; Table S7). Similarly, when compared to the sperm and seminal fluid proteins, we extracted 197 *B. anynana* spermatophore orthologs in *B. terrestris* (Table S7), 207 from *A. aegypti* (Table S7), and 431 from *D. melanogaster* (Table S7). Since the spermatophore samples were encased in the female bursa, we identified orthologs that were uniquely female derived proteins in *D. melanogaster* female reproductive tracts from [79]. This approach identified 22 proteins that could be of female origin in *B. anynana,* solely based on orthology (Table S8). Lastly, 35.5% (761) of *B. anynana* spermatophore proteins contained a predicted signal peptide sequence, suggesting their possible secretory functions (Table S9). These proteins belonged to serine protease related activity, proteolysis, and odorant binding GO terms.

### 4. Male experience increased DA proteins transferred to preferred and unpreferred females

Significantly differentially abundant (DA) proteins obtained after permutation tests for specific treatment comparisons indicate that the experienced males mated to 0 DHSN versus 2 DHSN female (E0sp vs E2sp) had the highest uniquely DA proteins out of our six comparisons (Figure 3A). We identified 232 DA proteins from the E0sp vs E2sp comparison (Figure 3B, Table S10), of which 161 were uniquely DA to this comparison. We recovered 117 DA proteins for E0sp vs N0sp (Figure S5A, Table S11); 102 DA proteins for E2sp vs N2sp (Figure S5B, Table S12); 108 DA proteins for N0sp vs N2sp (Figure S5C, Table S13); 99 DA proteins for 0 DHSN vs 2 DHSN (Figure 3C, Table S14), and 81 DA proteins for experienced vs naïve (Figure 3D, Table S15). We did not recover any enriched GO terms for any of these treatment combinations when compared to the spermatophore proteome as the reference set. However, when we set the reference set to the *Bicyclus anynana* genome, we recovered 62 enriched GO terms from the E0sp versus E2sp contrast (FDR<0.01; Table S16), 17 enriched GO terms from the 0 DHSN versus 2 DHSN contrast (FDR<0.01; Table S17), and 2 enriched GO terms from the experienced versus naïve contrast (FDR<0.01; Table S18). Some of the GO terms from the E0sp versus E2sp contrast include molecular functions such as “carbohydrate binding”, and “nucleotide binding” as well as biological processes such as “proteolysis”, and “protein catabolic process”. From the 0 DHSN versus 2 DHSN contrast, some of the enriched GO terms include peptide and amide “biosynthetic” and “metabolic process”. The two enriched GO terms from experienced versus naïve contrast both belong to “non-membrane-bounded organelle”. We did not recover any significantly enriched GO terms from the other three contrasts (E0sp versus N0sp; E2sp versus N2sp; and N0sp versus N2sp).

**Figure 3:**
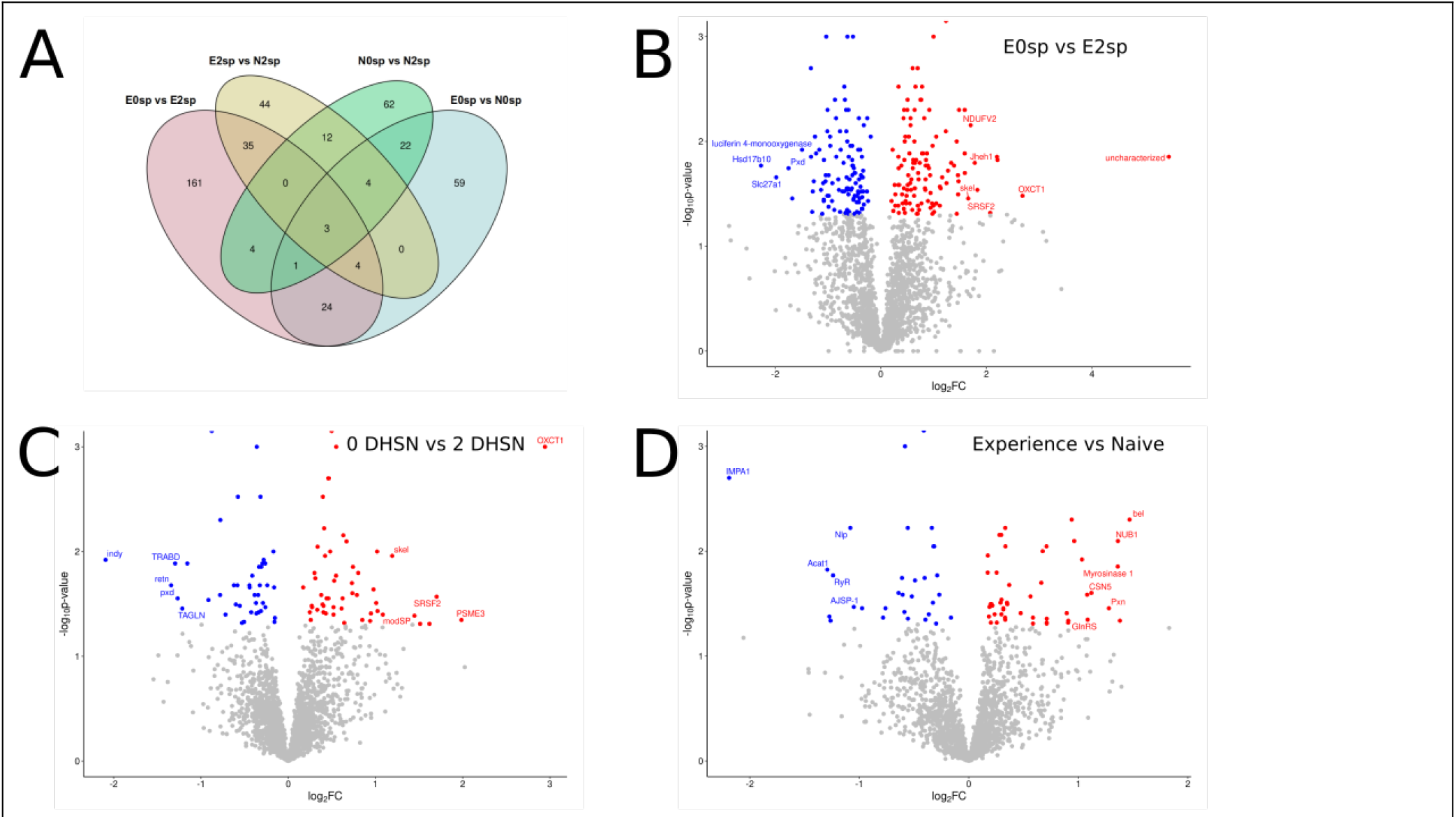
Spermatophore proteins change in response to previous male experience and males’ perception of female phenotype. A) A venn diagram depicting the number of common and unique differentially abundant (DA) proteins for a select contrasts. Experienced males mated to 0 DHSN or 2 DHSN females (E0sp vs E2sp) had the highest number of unique DA proteins. B) Volcano plot depicting DA proteins in E0sp vs E2sp contrast. Red dots indicate proteins that have over abundance in E0sp compared to E2sp treatment (p<0.05), whereas blue dots indicate proteins that have lower abundance in E0sp compared to E2sp treatment (p<0.05). Grey dots are proteins that are not DA (p>0.05). C) DA proteins in spermatophores transferred to 0 DHSN or 2 DHSN females, irrespective of male experience. D) DA proteins in spermatophores transferred by experienced versus naïve males, irrespective of female phenotype. E0sp= Experienced males mated to 0 DHSN females; E2sp= Experienced males mated to 2 DHSN females; N0sp= naïve male mated to 0 DHSN females; N2sp= naïve male mated to 2 DHSN females.

### 5. Proteins associated with oogenesis, neural signalling, learning and memory are DA across mating treatments

We found a number of DA proteins associated with oogenesis, neural signalling, learning and memory. Some proteins associated with oogenesis that are uniquely DA in 0 DHSN versus 2 DHSN treatment include *ATP-dependent RNA helicase vasa* (log2FC= -0.99, Figure 4I), and *transgelin* (log2FC= -1.21). Other proteins associated with oogenesis include *protein dead ringer isoform X2* which is DA in 0 DHSN or 2 DHSN contrast (log2FC= -1.34, Figure 4H), and in N0sp versus N2sp contrast (log2FC= -1.56), *profilin* which is DA in 0 DHSN versus 2 DHSN (logFC= 0.31), E0sp versus N0sp (log2FC= -0.30), E2sp versus N2sp (log2FC= -0.34), N0sp versus N2sp (log2FC=0.27), and experienced versus naïve contrast (log2FC= -0.33), and *COP9 signalosome complex subunit5* which is DA between spermatophores transferred by naïve and experienced males (log2FC=, Figure 4D). We also recovered proteins related to neural signalling, axon and dendritic growth, neurodevelopment, and learning and memory formation such as *protein enhancer of sevenless 2B* (E0sp vs E2sp log2FC= 0.56; E2sp vs N2sp log2FC= -0.57), *glutamine tRNA ligase* (Experienced vs naïve log2FC=; Figure 4E), *SRC kinase signalling inhibitor 1-like* ( Experienced vs naïve log2FC=; Figure 4F), and *glutamate dehydrogenase* (E0sp versus E2sp log2FC= -0.67; N0sp versus N2sp log2FC= -0.89; 0 DHSN or 2 DHSN log2 FC= -0.77). Proteins uniquely DA in E0sp versus E2sp contrast included *torsin 1A* (log2FC= -0.77), *renin receptor* (log2FC= 0.54), and *proton-coupled amino acid transporter-like protein pathetic* (log2FC= 0.41).

**Figure 4:**
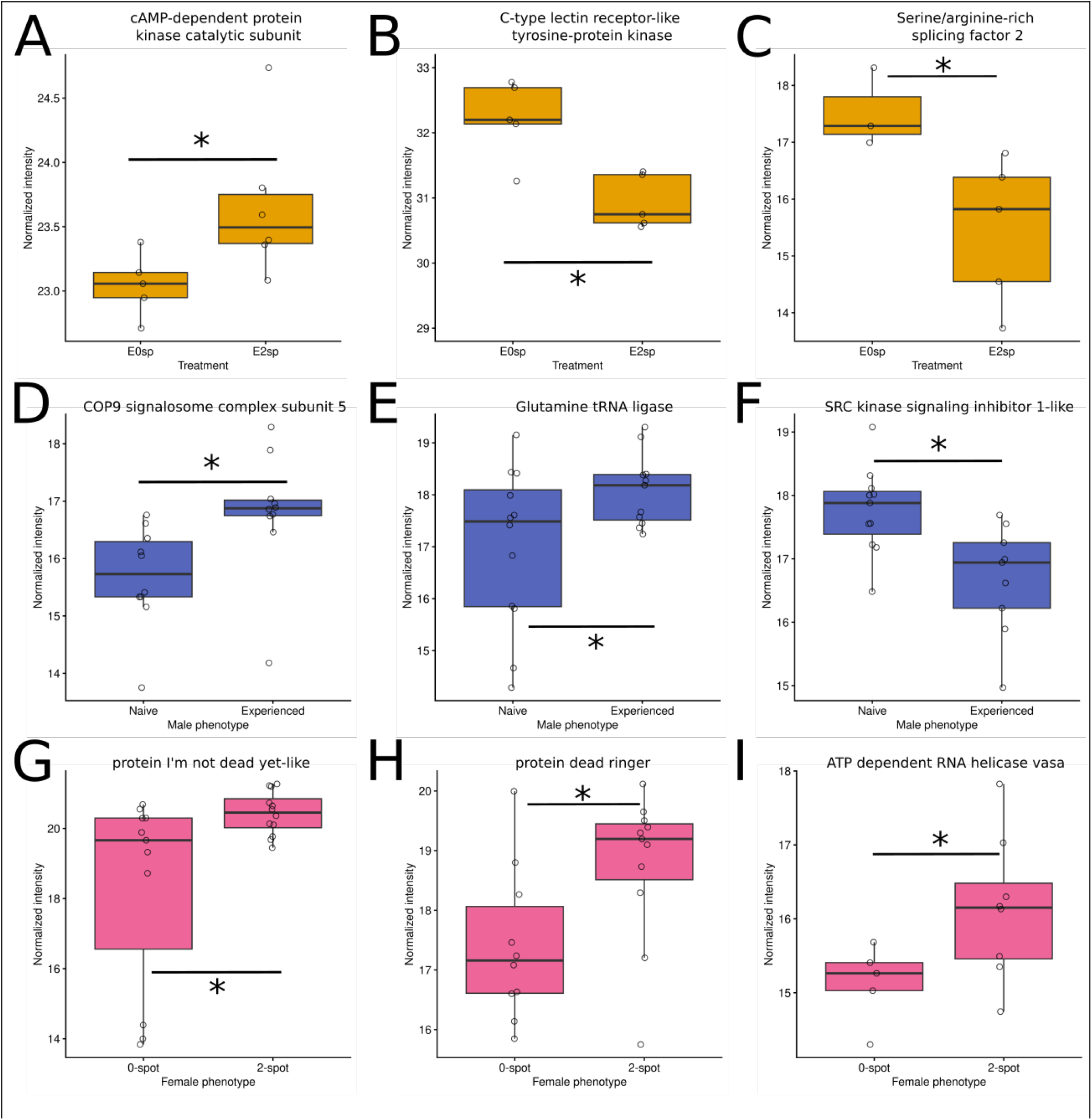
Proteins driving diverse biological processes are DA in spermatophores based on male mate preference learning, and female phenotype. Many proteins that are implicated in oogenesis, neural development, learning and memory formation, circadian rhythm, and female receptivity to re-mating are DA in spermatophore transferred by (A, B, C) experienced male mated either to 0 DHSN or 2 DHSN female (E0sp versus E2sp comparison), (D, E, F) experienced versus naïve male, and (G, H, I) proteins in spermatophore transferred to either 0 DHSN or 2 DHSN females. *p<0.05 after permutation tests.

We also recovered proteins related to circadian rhythm behaviours that were DA. *Protein quiver* (log2FC= 0.38) and *cAMP-dependent protein kinase catalytic subunit* (log2FC= -0.61, Figure 4A) were DA in our E0sp vs E2sp contrast, and while *protein takeout* which was DA in E0sp versus N0sp contrast (log2FC= -1.11). Other proteins that are implicated in various other processes were noteworthy. *Protein I’m not dead yet* (*Indy,* 0 DHSN versus 2 DHSN log2FC=, Figure 4G) is implicated in lifespan, *serine/arginine-rich splicing factor 2* (E0sp versus E2sp log2FC=; Figure 4C) is involved in mRNA processing, and *C-type lectin receptor-like tyrosine-protein kinase* (E0sp versus E2sp log2FC=; Figure 4B) is a known sex peptide network protein in *D. melanogaster* [80] transferred in the ejaculates during mating, but its function in our butterfly is unknown.

## Discussion

Here we described the proteomic changes in spermatophores transferred by males to females associated with male experience and perception of female phenotype in the butterfly *Bicyclus anynana*. To that end, we first characterized and described the spermatophore proteome of a wet season *B. anynana* where we recovered more than 400 enriched GO terms in various molecular functions, cellular components, and biological functions. We then found that without any changes related to courtship and mating related behaviours in one-choice assays, greater uniquely DA proteins were present between experienced males mated to 0 DHSN versus 2 DHSN females than between naïve males mated to 0 DHSN versus 2 DHSN females, with DA proteins associated with neural development, axon morphogenesis, learning, and circadian rhythm. Additionally, proteins related to oogenesis were DA between spermatophores transferred to 0 DHSN versus 2 DHSN females, when pooling across experienced versus naïve male conditions.

A main objective of this study was to characterize the spermatophore proteome of a wet season *Bicyclus anynana.* We identified 2144 proteins, and GO enrichment identified terms needed for the spermatophore’s main function i.e sperm motility and fertilization. The GO terms of the proteome were complementary to other lepidoptera [36–41,43,81] and insects more broadly [35,42,44,72,75]. Top most-specific enriched GO terms in *B. anynana* spermatophore proteome included ATP and GTP binding, and GTPase activity. These terms indicate high metabolic requirements and the need for energy during various cellular and molecular processes like post translational modifications of proteins, ion binding [46], and sperm motility in fish [82]. In line with that, we found the greatest number of proteins associated with the biological processes GO terms of protein folding and intracellular ion transport in the *B. anynana* spermatophore proteome. Over enrichment of the same GO terms (ATP binding, GTP binding, and GTPase activity) identified in the chicken (*Gallus gallus*) seminal plasma extracellular vesicles [48] indicate conserved molecular functions of those seminal fluid proteins across animals. Other enriched terms included GTP binding and GTPase activity, and proteins in these terms belonged to *Ras-related* and *Rab-related protein family,* associated with cell signalling and acrosome functioning [83,84], which control male fertility [85]. Many of these *Ras-related proteins* are also associated with neural functioning [86] and their abundance in the spermatophore proteome may indicate neural cell binding functions that, if bound to female neural cells, can influence post-mating female physiological and behavioural changes. Consequently, we also identified Rab-related signal transduction terms over enriched in the *B. anynana* spermatophore proteome, suggesting that *Ras* and *Rab-related* protein functions that rely on GTP binding and GTPase activity are important for intercellular communication (such as sperm-egg interactions mediated by the acrosomes) that directly control male fertility and sperm capacitation [87]. Other terms enriched in the *B. anynana* spermatophore proteome include the binding of calcium ion and actin. Actin regulates many nuclear and cellular modifications, and transport of materials during spermatogenesis whereas calcium ions are required for triggering sperm motility in rainbow trout fish [45] and in egg activation in females across animals [88–91]. However, whether calcium is transferred to females via male ejaculates in butterflies is unknown.

We found different GO terms common between *B. anynana* spermatophore proteome and other lepidoptera proteomes [36–40,43], possibly because of the cells used to generate those proteomes. When compared to the *Heliconius* spermatophore proteome, which included the whole spermatophore to generate the proteome [40], we found many “binding” terms common with the *B. anynana* spermatophore proteome. But when we compared the *B. anynana* to the *Danaus plexippus*, *Pieris rapae,* and *Manduca sexta* sperm proteome, which included only the sperm to generate the proteome [36,38,43], we found many purine associated GO terms to be common. This suggests that the seminal fluid proteins in the spermatophore may have many “binding” related functions (RNA binding, protein binding, calcium ion binding, metal ion binding), which may be useful in post-mating processes that influence female physiology and behaviour.

Without behavioural and spermatophore size differences between treatments in our one-choice trials, we found the greatest difference in proteins given to 0 DHSN (preferred) females versus 2 DHSN (unpreferred) females by experienced males. This difference in proteins corresponds to previously observed learned preferences in this species, where trained *B. anynana* males prefer to mate with 0 DHSN over 2 DHSN females in two-choice assays, while naïve males mate randomly [58]. Thus, our findings suggest males are able to tailor their spermatophore proteins based on female phenotype [22] after acquiring a preference. We found many circadian proteins to be DA, which have previously been implicated in changing female circadian rhythms and rhythms of other genes and proteins in the mated female [33]. Two such proteins are *cAMP-dependent protein kinase catalytic subunit* that induces light-dependent protein kinase activity in brains [92] controlling learning and memory formation, and *protein quiver* which is required for sleep homeostasis and after sleep deprivation [93]. Lack of sleep can affect reproductive traits such as egg laying in females [94]. Thus, increased abundance of circadian proteins towards preferred females (in case of *quiver*) may increase fecundity and reproductive fitness of the male, based on his acquired preference through learning. Mating has also been shown to increase feeding in some female insects [27,28], as well as intake of materials from the spermatophore itself (as spermatophores are nuptial gifts), for which binding of food is an important process after mating. We found over enrichment of carbohydrate binding proteins, nucleotide binding proteins, and many lipid binding proteins to be DA in the spermatophores given by experienced males to preferred and unpreferred females. Mating also induces gut growth and expansion [95–97], possibly for food and nutrient absorption, and increased absorption of food and spermatophore materials provides energy for reproductive physiological and behavioural processes such as oogenesis, ovulation, and egg laying [95,96]. Future studies should examine whether these proteins transferred to females during copulation do influence female circadian behaviours in *B. anynana*, and consequently reproductive output and male fecundity.

We recovered many oogenesis and ovulation proteins DA in spermatophores given to 0 DHSN and 2 DHSN females, irrespective of male experience (naïve or experienced). These proteins could be responsible for more eggs laid by 0 DHSN females [59]. Mutations in protein *I’m not dead yet* (*indy)* in *Drosophila melanogaster* increase the lifespan of flies [98], and lower expression of *indy* increases fecundity [99]. *Indy* had lower abundance in spermatophores given to 0 DHSN females compared to 2 DHSN females, and this protein could be associated with the previously observed increased egg laying in 0 DHSN *B. anynana* females [59]. Other oogenesis proteins that had lower abundance in the spermatophores provided to 0 DHSN females include *protein dead ringer* (*retn*), which is involved in oogenesis [100], female receptivity, and male courtship behaviours [101] and *RNA helicase vasa,* which is a nucleic acid and ATP binding protein required during early oogenesis [102]. Lastly, *protein profilin* was abundant in spermatophores transferred to 0 DHSN females, which is involved in the transport of intercellular cytoplasm during oogenesis [100], as well as regulating actin during neuron projection, neurogenesis, and brain development [103]. Overall, many oogenesis proteins as well as neural proteins were found to be DA in spermatophores transferred to 0 DHSN female and 2 DHSN female, suggesting that these proteins could alter the observed difference in egg laying by *B. anynana* females [59], and could also control female behaviour, thereby alleviating the realized fitness of males based on their acquired (through learning) or cryptic innate preference. Future research should explore the functional role of these transferred proteins in influencing female behaviour, lifespan, and reproductive output.

## Conclusions

In this study, we set out to explore the consequence of male preference acquisition through learning by exploring the spermatophore proteins transferred by said males to their preferred and unpreferred females. Subsequently, we characterized the first spermatophore proteome of the wild type wet season lab reared *B. anynana,* revealing 2144 unique proteins belonging to more than 400 GO term categories. The spermatophore proteome is comparable with other lepidoptera spermatophore and sperm proteomes. Next, we found that without any changes in pre- and peri-copulatory behavioural traits and spermatophore size, we identified differentially abundant spermatophore proteins based on male social experience (naïve and experienced) and the males’ perception of the female phenotype (0 UV DHSN and 2 UV DHSN female), suggesting that male butterflies are able to plastically alter what they invest in their spermatophore based on their past and present social environmental cues. These changes in proteins were largely reflected in experienced males mating with either their preferred (0 DHSN) or unpreferred (2 DHSN) females. Proteins that influence many traits such as oogenesis, circadian rhythm, neural signalling, binding, and sperm-egg interactions were differentially abundant across treatments, and many oogenesis proteins had lower abundance in male spermatophores transferred to 0 DHSN females compared to 2 DHSN females, irrespective of male social experience. Taken together, our study characterizes the consequence of a learning acquired pre-mating preference on peri- and post-mating transfer of spermatophore proteins. It highlights the roles of spermatophore proteins in influencing female physiology and behaviours that are vital for the pair’s reproductive fitness, that governs the evolution of traits and preferences via post-copulatory sexual selection.

## Ethics

All butterflies were reared in a climate controlled greenhouse set at ∼27°C with 13:11 L:D cycle and 70-80% RH, which are similar conditions experienced in their natural habitat, as stated in the U.S. Department of Agriculture, Animal and Plant Health Inspection Service permits P526P-17–00343 and P526P20–00417. All adult butterflies were fed with ad libitum wet banana everyday, and only individuals that were used for the experiment were sacrificed.

## Data accessibility

Raw behavioural data is available in Supplementary Table 1. The MS proteomics data have been deposited on ProteomeXchange Consortium via PRIDE, and can be accessed using the identifier PXD081208. All scripts are available on Github and can be accessed using https://github.com/sdpotdar/Proteomic-consequences-of-male-mate-preference-learning-and-female-phenotype-in-Bicyclus-anynana.

## Declaration of AI use

We declare that we took assistance from an AI-powered search engine (Perplexity.ai) in refining a part of our bioinformatics custom scripts. All scripts were manually checked for accuracy. We did not use AI for any other part during the preparation of this manuscript.

## Author contributions

SP-conceptualization, methodology, investigation, formal analysis, visualization, writing-original draft, writing-review and editing; DP-formal analysis, funding acquisition, supervision, writing-original draft, writing-review and editing, ELW-conceptualization, methodology, funding acquisition, supervision, visualization, writing-original draft, writing-review and editing.

## Acknowledgments

We thank Grace Hirzel, Matthew Murphy, David A. Ernst, Yi Ting Ter, Kiana Kasmaii, and Keity Farfán Pira for their assistance in butterfly husbandry. We thank Jeffrey A. Lewis, Brian A. Counterman, William J. Etges, Nathan Morehouse, and James Walters for their helpful suggestions and discussions during this project. We also thank Alan Tackett, Rick Edmondson, Sam Mackintosh, Lisa Orr, Robert Brown, and Stephanie Byrum from IDeA National Resource for Quantitative Proteomics at UAMS Little Rock for their help in spermatophore protein sample preparations and analysis. We thank Aarna Kulkarni for re-analyzing spermatophore areas, Crystal Crook for discussions and insights on differential protein analysis, and Jolie A. Carlisle for orthology analysis, and reviewing the manuscript.

## Supplementary Figures

**Figure S1:**
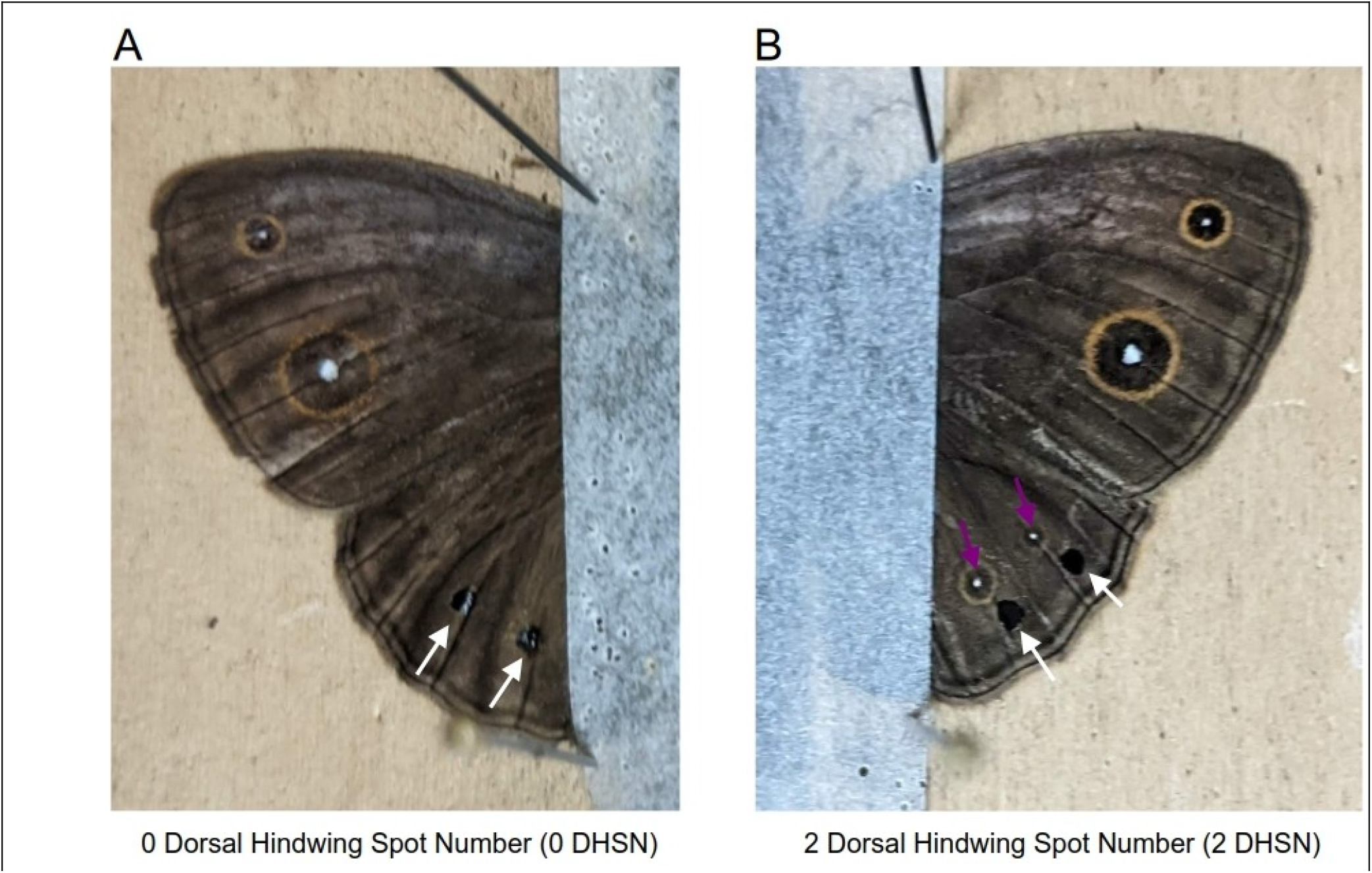
Visual depiction of 0 and 2 DHSN females: The dorsal wing surfaces of a female *B. anynana* with (A) naturally occurring UV spots on the dorsal hindwing blocked by black paint (white arrows in A) making them a 0 DHSN female, while (B) the black paint is applied just below (white arrows in B) the two naturally occurring dorsal hindwing UV spots (purple arrows in B), making them a 2 DHSN female.

**Figure S2:**
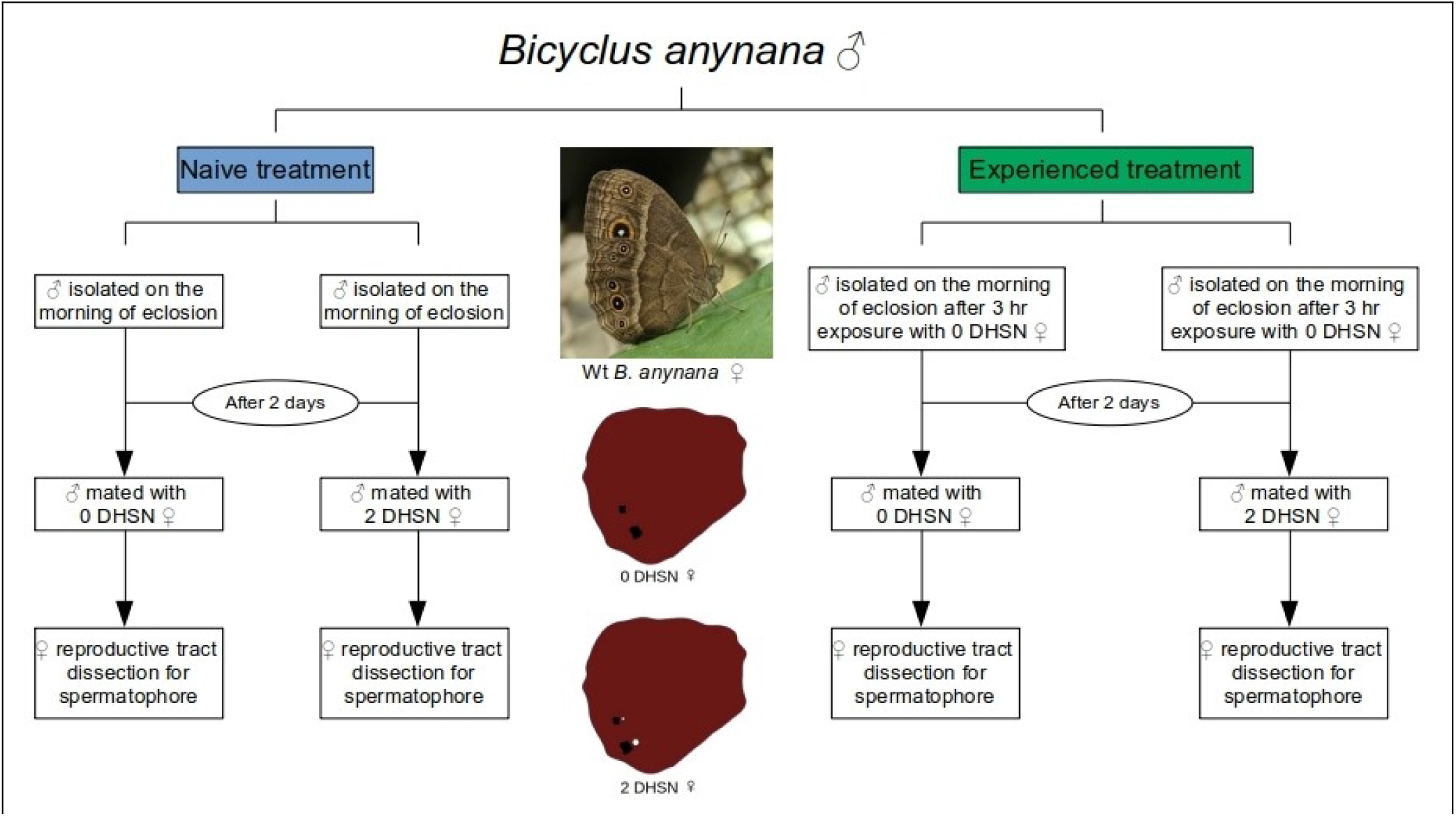
Experimental design: Experimental design to assess spermatophore proteins transferred by naïve and experienced males to their preferred (0 DHSN) or unpreferred (2 DHSN) females. naïve males were isolated after eclosion and given either a 0 DHSN or a 2 DHSN female to mate on day 2. Experienced males were exposed to a 0 DHSN female for 3 hours on the day of eclosion and isolated. After two days (day 2), experienced males were either given a 0 DHSN or a 2 DHSN female to mate. Successfully mated pairs were frozen and male transferred spermatophores were dissected from the female reproductive tract for downstream spermatophore proteomic analyses. All females were naturally 2 DHSN, and 0 DHSN females were generated by blocking the UV spots with a black paint. 2 DHSN females were also painted with black paint just below the naturally existing UV spots (See Figure S1).

**Figure S3:**
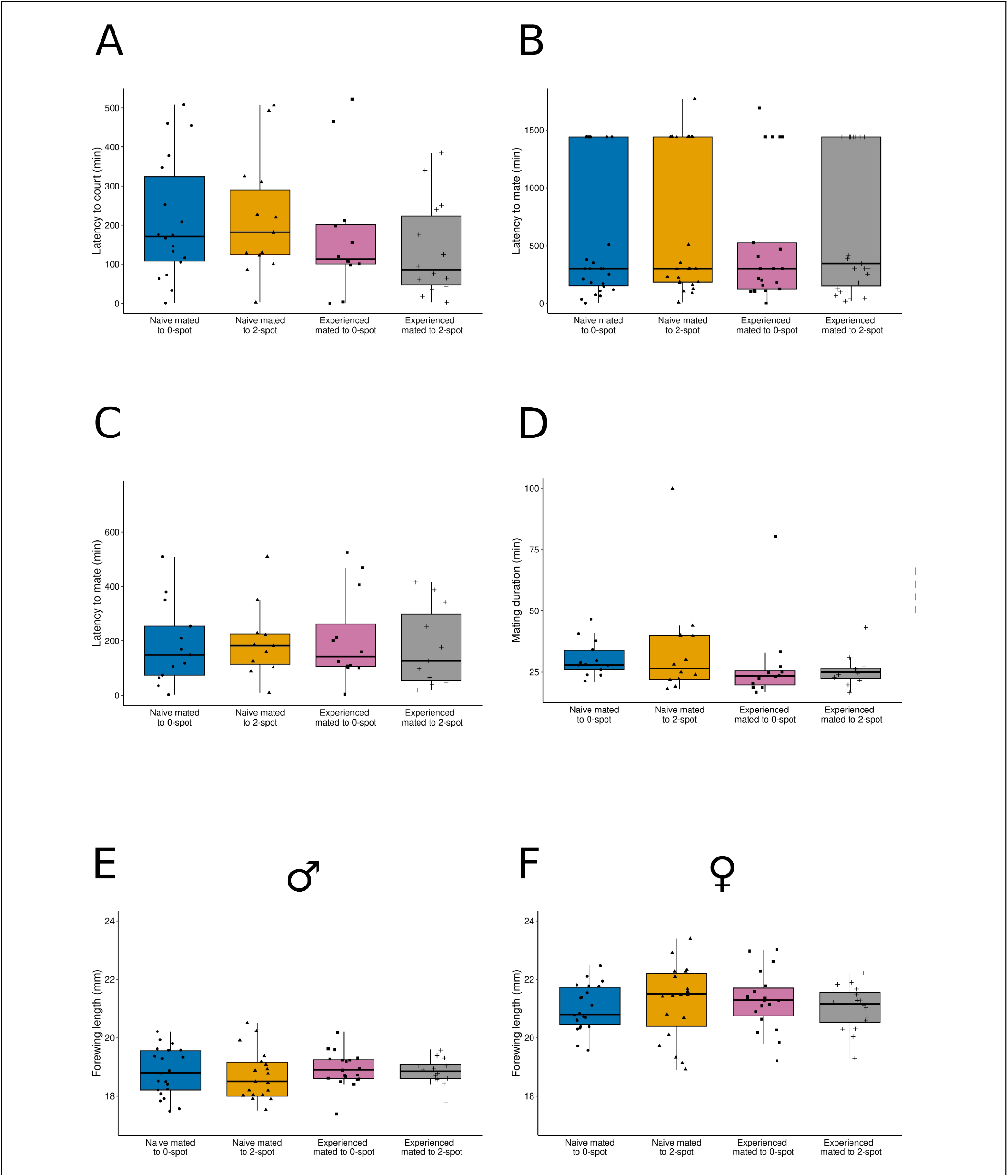
Mating related behaviours and butterfly size did not change across treatments. The males’ A) latency to court, B) latency to mate (including pairs that did not mate), C) latency to mate (excluding pairs that did not mate), D) mating duration, E) male fore wing length as a proxy for size, and F) female fore wing length as a proxy for size did not change across the four different treatments (naïve mated to 0 DHSN, naïve mated to 2 DHSN, Experienced mated to 0 DHSN, Experienced mated to 2 DHSN).

**Figure S4:**
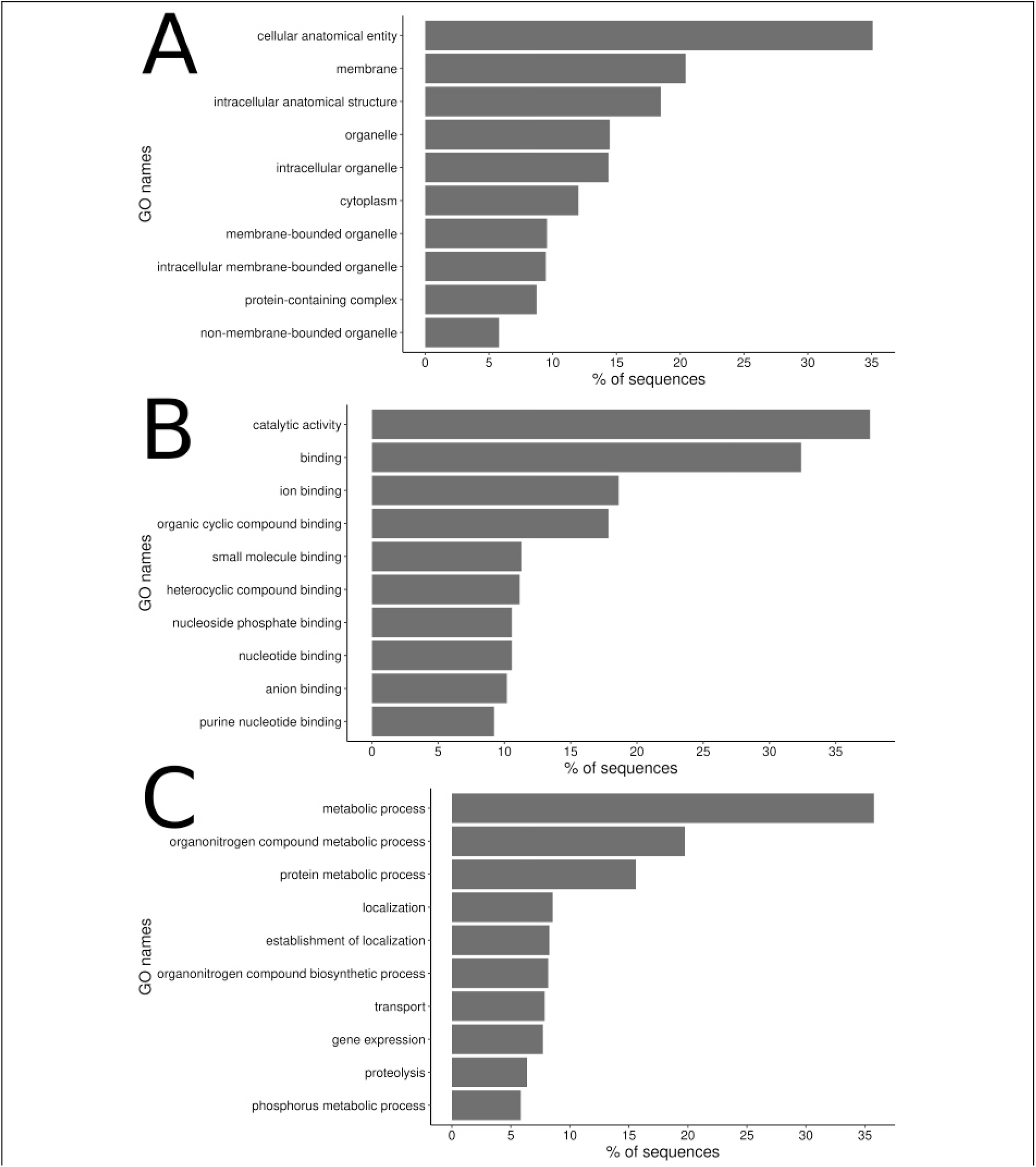
Top enriched GO terms in the *B. anynana* spermatophore proteome. Top enriched GO terms in the *B. anynana* spermatophore proteome in the A) cellular component category, B) molecular function category, C) biological processes category

**Figure S5:**
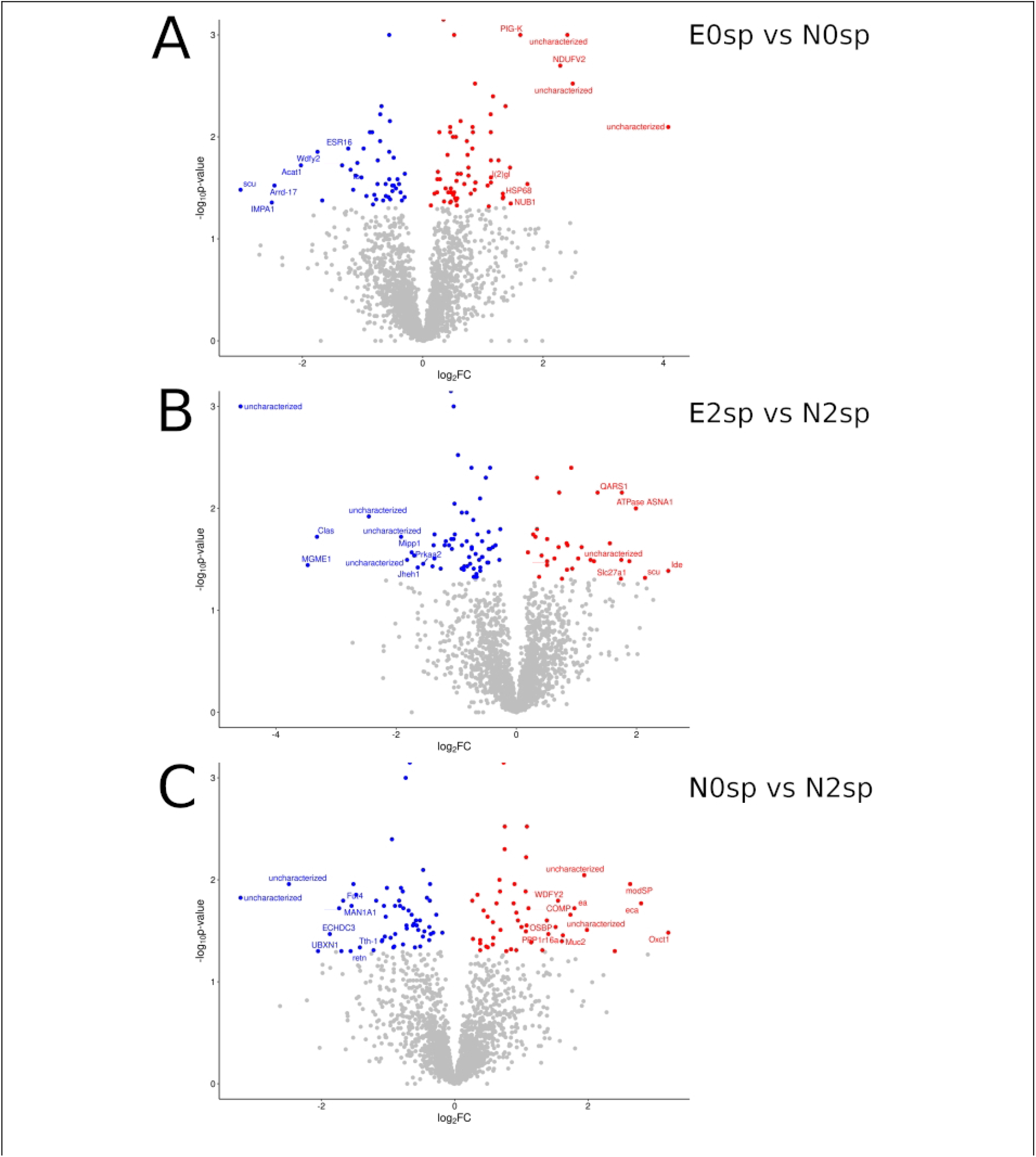
Male experience and female phenotype affect differential abundance in proteins in A) E0sp vs E2sp contrast, B) E2sp vs N2sp contrast, C) N0sp vs N2sp contrast. Red dots indicate proteins that have over abundance in the term that appears first in a contrast (example E0sp in E0sp vs E2sp contrast) (p<0.05), whereas blue dots indicate proteins that have lower abundance in the term that appears first in the contrast (p<0.05). Grey dots are proteins that are not DA (p>0.05).

